# Sulfated RaxX, which represents an unclassified group of ribosomally synthesized post-translationally modified peptides, binds a host immune receptor

**DOI:** 10.1101/442517

**Authors:** Dee Dee Luu, Anna Joe, Yan Chen, Katarzyna Parys, Ofir Bahar, Rory Pruitt, Leanne Jade G. Chen, Christopher J. Petzold, Kelsey Long, Clifford Adamchak, Valley Stewart, Youssef Belkhadir, Pamela C. Ronald

## Abstract

The rice immune receptor XA21 is activated by the sulfated microbial peptide RaxX (required for <u>a</u>ctivation of <u>X</u>A21-mediated immunity <u>X</u>) produced by *Xanthomonas oryzae* pv. *oryzae* (*Xoo*). Mutational studies and targeted proteomics revealed that RaxX is processed and secreted by the protease/transporter RaxB, whose function can be partially fulfilled by a noncognate <u>p</u>eptidase-<u>c</u>ontaining transporter <u>B</u> (PctB). RaxX is cleaved at a Gly-Gly motif, yielding a mature peptide that retains the necessary elements for RaxX function as an immunogen and host peptide hormone mimic. These results indicate that RaxX is a founding member of a previously unclassified and understudied group of tyrosine sulfated RiPPs (<u>ri</u>bosomally synthesized, <u>p</u>ost-translationally modified <u>p</u>eptides). We further demonstrate that sulfated RaxX directly binds XA21 with high affinity. This work reveals a complete, previously uncharacterized biological process: bacterial RiPP biosynthesis, secretion, binding to a eukaryotic receptor and triggering of a robust host immune response.

## INTRODUCTION

<u>Ri</u>bosomally synthesized and <u>p</u>ost-translationally modified <u>p</u>eptides (RiPPs) include anti-microbial, anti-cancer, insecticidal, and quorum sensing peptides (Arnison et al., 2013). RiPPs are structurally and functionally diverse yet share commonalities. They are ribosomally synthesized as a precursor peptide with a cleavable N-terminal leader and a post-translationally modified core that becomes the final secreted bioactive RiPP (Figure 1A; Arnison et al., 2013). Many RiPP leaders direct the modification and/or export proteins to the core. This permits the biosynthetic proteins to act on a diverse range of core peptides while maintaining substrate specificity (Oman and van der Donk, 2010) and allows for leader peptide-guided designed synthesis of hybrid RiPPs (Burkhart et al., 2017).

**Figure 1.**
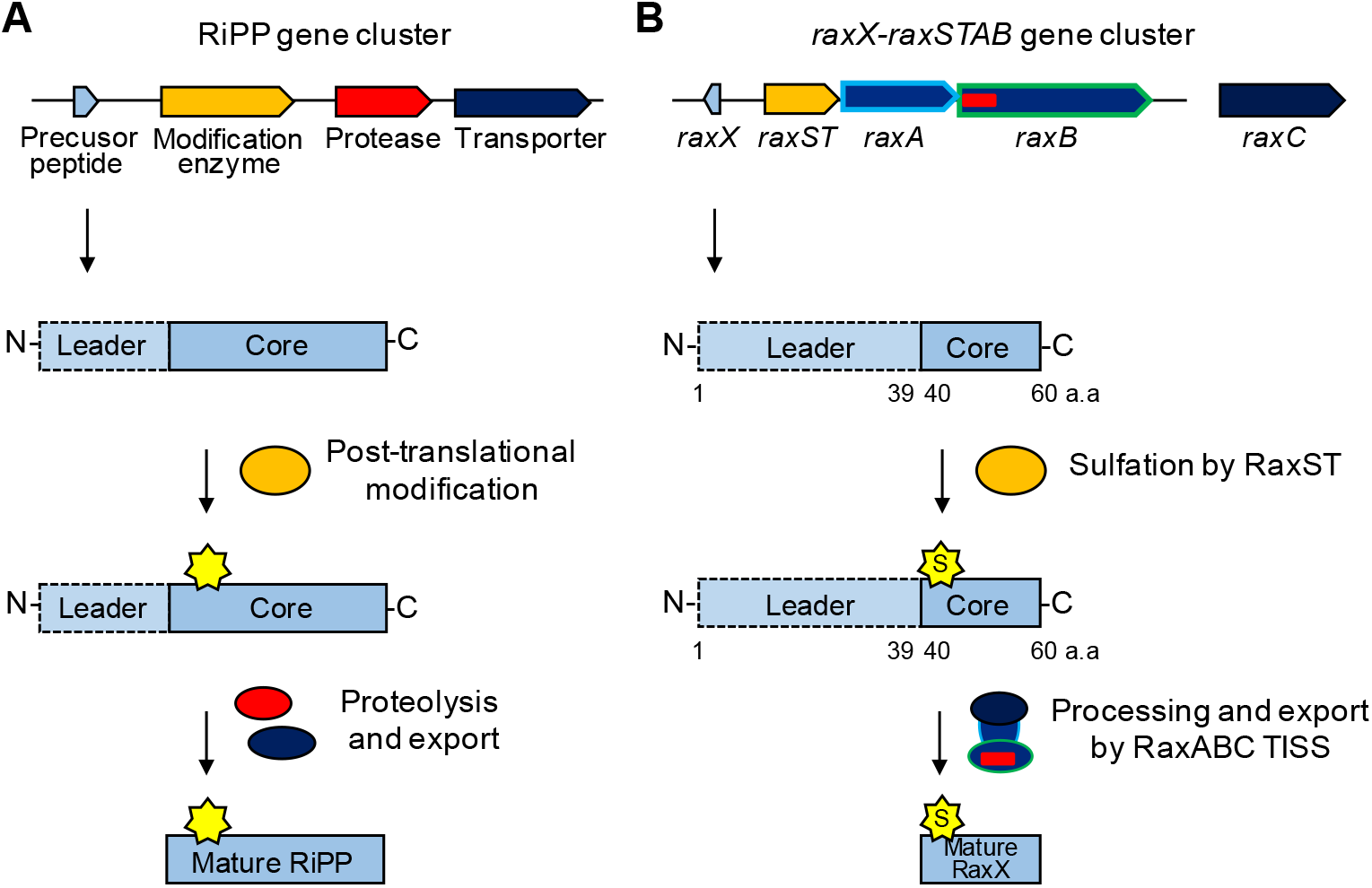
Biosynthetic pathways of RiPPs and RaxX. (A) General RiPP biosynthetic pathway. The RiPP precursor and biosynthetic proteins are ribosomally synthesized. The core, which becomes the final RiPP product, is post-translationally modified by enzyme(s) encoded in the same genomic region. The N-terminal leader is enzymatically removed by a protease and the modified core is exported by a transporter, releasing the mature bioactive RiPP. (B) RaxX biosynthetic pathway. The RaxX precursor is ribosomally synthesized and the core is sulfated by the sulfotranferase RaxST encoded upstream. We hypothesize that the peptidase-containing ABC (ATP-binding cassette) transporter (PCAT) RaxB removes the N-terminal leader and transports the sulfated mature RaxX peptide through the type I secretion system (T1SS) composed of RaxB, the periplasmic adaptor protein RaxA, and the genetically unlinked outer membrane protein RaxC.

RiPPs achieve substantial chemical diversity through extensive post-translational modifications and are divided into over 20 groups (Arnison et al., 2013). One group that has not been well-studied nor formally categorized as RiPPs are tyrosine sulfated peptides.

Tyrosine sulfation is a post-translational modification that influences receptor-ligand binding in diverse host-microbe interactions (Stone et al., 2009). For example, in humans, tyrosine sulfation of the integral membrane protein CCR5 (C-C chemokine receptor type 5) significantly enhances binding of the human immunodeficiency virus (HIV) envelope glycoprotein, facilitating HIV entry (Farzan et al., 1999).

In rice, the transmembrane immune receptor XA21, which shares similarities to animal TOLL like receptors (TLRs) and the *Arabidopsis* flagellin-sensitive 2 (FLS2) and EF-Tu receptors (EFR) (Dardick et al., 2012), responds to sulfated derivatives of the microbial peptide RaxX (<u>r</u>equired for <u>a</u>ctivation of <u>X</u>A21-mediated immunity <u>X</u>) produced by the Gram-negative pathogenic bacterium *Xanthomonas oryzae* pv. *oryzae* (*Xoo*) (Pruitt et al., 2015, 2017; Schwessinger et al., 2016; Wei et al., 2016). Tyrosine sulfated RaxX16, a 16-residue synthetic peptide derived from residues 40-55 of the 60-residue RaxX precursor, is the shortest characterized immunogenic derivative of RaxX (Pruitt et al., 2017). RaxX16 shares a 13-residue sequence with high similarity to the 18-residue plant peptide hormone PSY1 (<u>p</u>lant peptide containing <u>s</u>ulfated t<u>y</u>rosine; Pruitt et al., 2017). PSY1 promotes cellular proliferation and expansion *in planta* (Amano et al., 2007) and increases root growth in both *Arabidopsis* and rice (Pruitt et al., 2017). Exogenous application of sulfated RaxX16 also promotes root growth (Pruitt et al., 2017), which is consistent with the hypothesis that RaxX mimics host PSY1 and PSY1-like proteins. These observations indicate that RaxX16 contains the biologically active domain of RaxX.

In *Arabidopsis*, PSY1 is ribosomally synthesized as a 75-residue precursor peptide and proteolytically processed into the mature secreted 18-residue sulfated and glycosylated peptide (Amano et al., 2007). Based on similarities between PSY1 and RaxX16, we hypothesized that sulfated RaxX is also processed and secreted to produce an extracellular molecule that can interact with host receptors.

The *raxX* gene is adjacent to the *raxSTAB* operon, encoding the RaxST tyrosylprotein sulfotransferase, which catalyzes sulfation of RaxX tyrosine residue 41, and two components of a predicted type I secretion system (T1SS): the RaxA periplasmic adaptor protein and the RaxB <u>p</u>eptidase-<u>c</u>ontaining <u>A</u>BC (ATP-binding cassette) transporter (PCAT; Figure 1B; Pruitt et al., 2015; da Silva et al., 2004). In other characterized PCATs, the N-terminal peptidase domain cleaves the substrate leader immediately after a Gly-Gly (GG) motif, and the processed substrate is secreted through the PCAT, periplasmic adaptor protein and TolC, an outer membrane channel protein (Kanonenberg et al., 2013).

Previously, we reported that derivatives of the immunogenic *Xoo* strain PXO99 containing null alleles of *raxX*, *raxST*, or *raxC* (encoding the *Xoo* TolC ortholog) evade XA21-mediated recognition, but strains with null alleles of *raxA* or *raxB* elicit a partial XA21-mediated immune response (Pruitt et al., 2015; da Silva et al., 2004). These results suggest that RaxA and RaxB are important for RaxX secretion but RaxX can also be released outside the cell independent of *raxA*/*raxB*.

Here, we report that synthetic RaxX peptide directly binds the extracellular domain (ECD) of the XA21 immune receptor (XA21^ECD^), an interaction enhanced by tyrosine sulfation of RaxX. We also demonstrate that RaxB is required for proteolytic processing and secretion of RaxX and identify a second PCAT PctB (<u>p</u>eptidase-<u>c</u>ontaining transporter component <u>B</u>) that can partially compensate for deletion of *raxB*. Our genetics and targeted proteomics data indicate that RaxX is ribosomally synthesized as a precursor peptide and cleaved downstream of a GG-motif in a RaxB-dependent manner. This proteolytic event releases the mature RaxX core peptide containing the sulfated Tyr-41 and the minimal 16-residue active region (Figure 1B). These results indicate that RaxX is a tyrosine sulfated RiPP, a previously undescribed class of RiPPs that mediate intercellular interactions.

## RESULTS

### Sulfated RaxX peptide binds XA21 with high affinity

We previously demonstrated that *in vivo* tyrosine sulfation of RaxX by RaxST is required to activate XA21-dependent immune responses (Pruitt et al., 2015), suggesting that sulfated, but not non-sulfated, RaxX is recognized by XA21. To test if RaxX directly binds to the XA21 immune receptor, we used microscale thermophoresis (MST), an optical tool that analyzes protein and small-molecule interactions (Wienken et al., 2010). The MST assays were conducted using XA21^ECD^ (XA21 residues 23-649) with varying concentrations of peptide to measure the dissociation constant (K_D_). When we titrated a synthetic 21-residue derivative of RaxX (RaxX21; residues 35-55) in the presence of XA21^ECD^, we observed a K_D_ of 400 nM with the non-sulfated form (RaxX21-nY; Figure 2). This K_D_ is 20-fold higher than the sulfated form (RaxX21-sY), which had a K_D_ of 20 nM (Figure 2), indicating that sulfated RaxX binds XA21 with higher affinity than non-sulfated RaxX.

**Figure 2.**
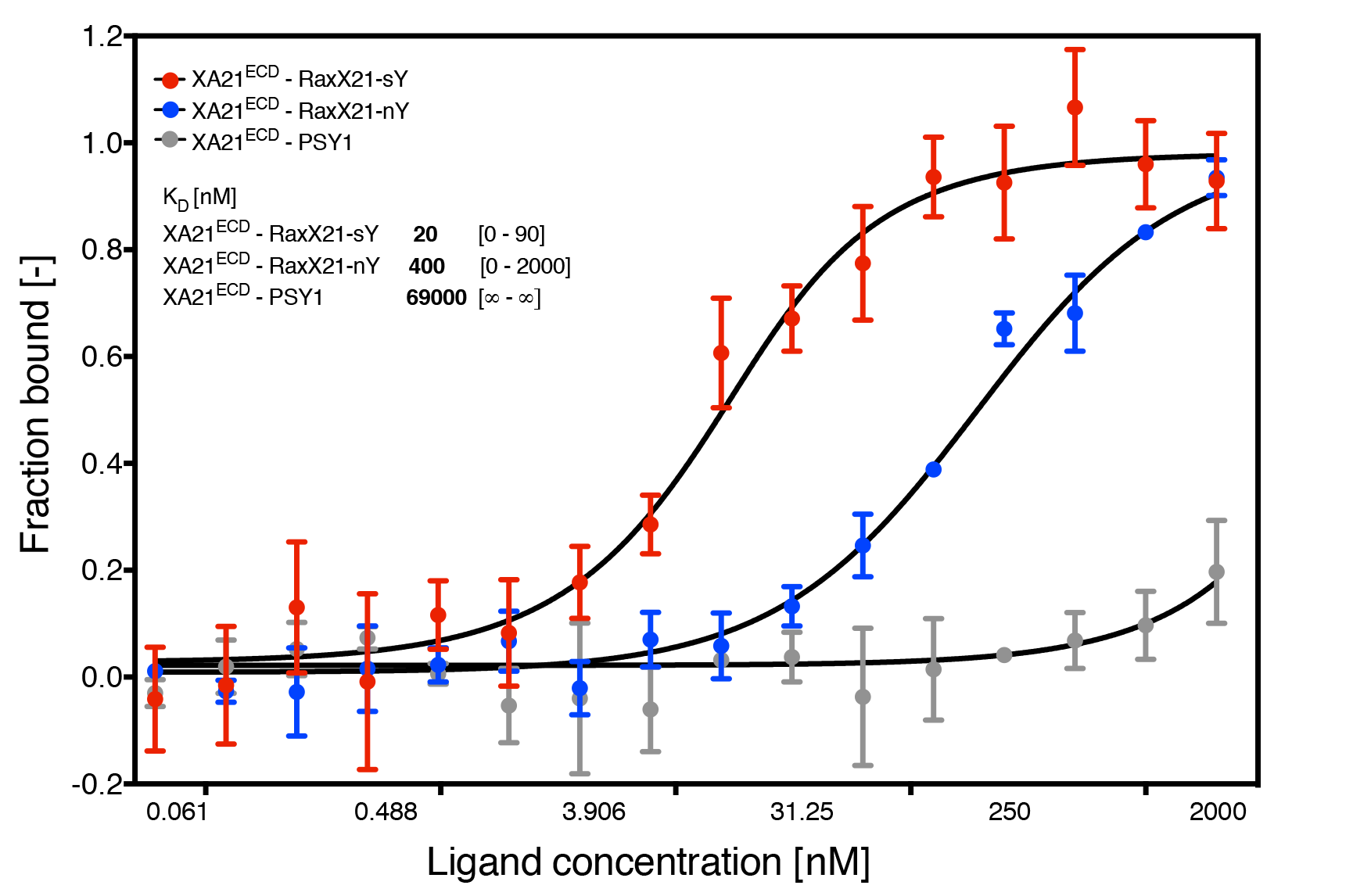
Sulfated RaxX peptide binds XA21ECD with high affinity. Quantification of binding between sulfated (-sY, in red) and non-sulfated (-nY, in blue) RaxX21 and sulfated PSY1 (in grey) peptides with the XA21 ectodomain (XA21^ECD^) by microscale thermophoresis (MST). Data points indicate the fraction of fluorescently labelled XA21 bound to the peptides during the assay (Fraction Bound [-]). The K_D_ and confidence intervals shown in brackets are indicated in nM. SE bars are representative of at least 2 independent measurements performed with independent protein preparations.

Although RaxX shares similarities with PSY1 in both sequence and root growth-promoting activities, synthetic sulfated PSY1 peptides derived from *Arabidopsis* and rice fail to activate XA21-dependent immune responses (Pruitt et al., 2017). Consistent with these reports, sulfated PSY1 failed to specifically bind XA21^ECD^ (Figure 2). These results indicate that RaxX binding to XA21^ECD^ is a specific interaction, which is enhanced by sulfation.

### SRM-MS detects RaxX as a secreted mature peptide

Based on its similarity to PSY1 and its direct binding to XA21^ECD^, we hypothesized that sulfated RaxX interacts with host receptors as a secreted proteolytically processed, mature peptide. To test this hypothesis, we assessed the presence of secreted RaxX in the extracellular milieu. Secreted proteins were harvested from *Xoo* strains grown in plant-mimicking media, concentrated, and digested with trypsin. The predicted mature and non-processed precursor RaxX peptides (represented by the DYPPPGANPK and HVGGGDYPPPGANPK tryptic peptides, respectively) were specifically detected by selected reaction monitoring-mass spectrometry (SRM-MS), which was calibrated using tryptic digests of RaxX16 (DYPPPGANPKHDPPPR) synthetic peptide and purified recombinant full-length RaxX (RaxX60; Figures 3A, S1A, and S1B).

**Figure 3.**
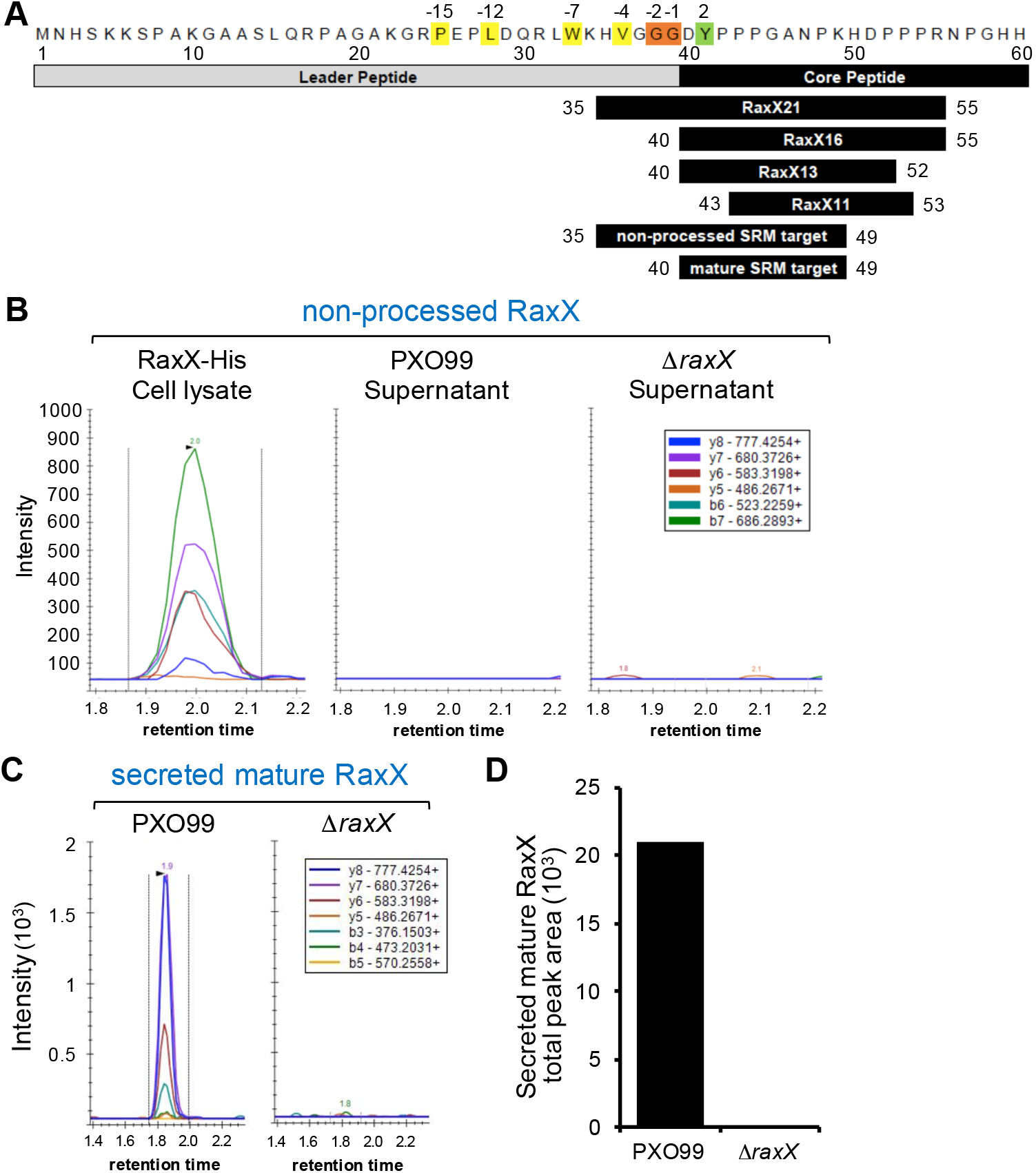
Mature RaxX is detected in *Xoo* supernatants. (A) RaxX precursor sequence. Numbers above refer to the residue position relative to the predicted cleavage site; numbers below relative to the precursor peptide. Cleavage of RaxX after Gly-38, Gly-39 (in orange) would result in a leader with hydrophobic residues (in yellow) at conserved locations typical of PCAT substrates and a core containing Tyr-41 (in green) sulfated by RaxST. Shown below this precursor are the synthetic RaxX peptide derivatives (RaxX21, RaxX16, RaxX13, and RaxX11) and the two tryptic peptide targets detected by selected reaction monitoring-mass spectrometry (SRM-MS). (B-C) SRM-MS chromatograms of the non-processed precursor (B) and predicted mature (C) RaxX tryptic peptides detected from Δ*raxX*(p*raxX-His*) cell lysates or from PXO99 or Δ*raxX* supernatants. Lines correspond to individual SRM transitions monitored. Legend indicates the detected peptide y-series fragment ion. (D) Quantification of the total peak area shown in C. See also Figure S1.

Similar to previous reports, we detected the RaxX non-processed precursor in the cell lysate (Figure 3B; Pruitt et al., 2015). However, only mature, processed RaxX was detected in the supernatant of the wildtype PXO99 strain (Figure 3). Mature RaxX was not detected in the Δ*raxX* negative control (Figures 3C and 3D), indicating that the observed peaks are specific for RaxX. Because trypsin is not known to cleave after glycine, which precedes the detected DYPPPGANPK tryptic peptide, these results suggest that the RaxX precursor is cleaved by an endogenous *Xoo* protease after Gly-39 and secreted outside the cell as a mature peptide (Figure 1B).

We also attempted immunoblot analysis as a complementary method to detect RaxX. However, an antibody generated against RaxX failed to detect RaxX16 or RaxX from *Xoo* supernatants (Figures S1C and S1D). We therefore used SRM-MS alone in subsequent experiments to verify RaxX secretion.

### Sequence analysis identifies a candidate secondary RaxX maturation and secretion system: PctAB

Prokaryotic RiPPs typically encode the maturation and secretion proteins in the same genomic region as the peptide (Figure 1A). We previously demonstrated that a strain carrying a null allele of *raxC*, encoding the predicted T1SS outer membrane protein, fails to activate XA21-mediated immunity, supporting a role for *raxC* in RaxX secretion (da Silva et al., 2004). In contrast, strains carrying null alleles of *raxB* or *raxA*, encoding the putative RaxX-associated PCAT and periplasmic adaptor proteins, respectively, maintain the ability to partially activate XA21-mediated immunity (da Silva et al., 2004). These results suggest that RaxX can be released in a *raxA*/*raxB*-independent manner.

The RaxB sequence contains an N-terminal C39 peptidase domain characteristic of PCATs (e.g. *Escherichia coli* colicin V (ColV) transporter CvaB), which have dual functionality as both a transporter and protease (Figure 4A; da Silva et al., 2004; Håvarstein et al., 1995; Wu and Tai, 2004). Query of the RaxB C39 peptidase domain sequence in BLAST searches of the PXO99 genome identified only one other candidate PCAT, which shares 50% sequence identity with RaxB (41% identity with *E. coli* CvaB). We named this gene (PXO_RS14825) as *pctB* (Figure 4B).

**Figure 4.**
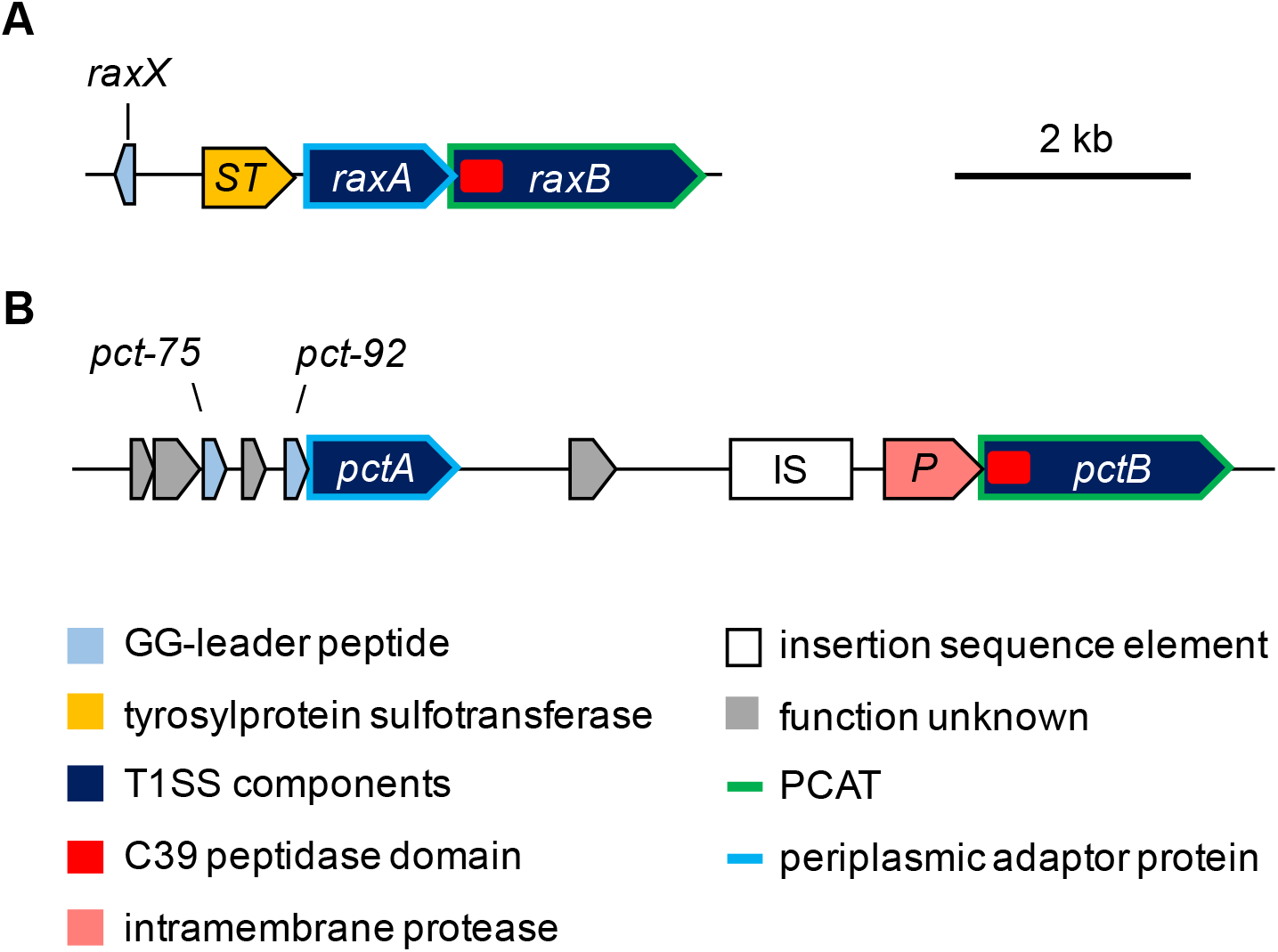
Genetic map of candidate RaxX biosynthetic proteins. (A) *raxX* is encoded upstream of the *raxSTAB* operon, containing genes for the RaxST sulfotransferase and components of a T1SS: the RaxA periplasmic adaptor protein and the RaxB PCAT. (B) Genes for components of a second peptidase-containing transporter system are not genetically linked to *raxX*: *pctA* and *pctB*, encoding a periplasmic adaptor protein and PCAT, respectively.

Immediately upstream of *pctB* is a gene, which we named *pctP* (PXO_RS14830), encoding a putative rhomboid family intramembrane serine protease (Figure 4B) that typically cleaves the transmembrane domain of membrane proteins (Freeman, 2014). Further upstream are an insertion sequence (IS*1112*), a region of uncertain heritage likely derived from multiple insertion and deletion events, and a predicted gene (PXO_RS14840), which we named *pctA*, encoding a putative periplasmic adaptor protein (Figure 4B). PctA shares 32% sequence identity with RaxA (24% identity with *E. coli* CvaA, the adaptor component of the ColV secretion system (Zhang et al., 1995)). Given the close proximity of *pctA* and *pctB* and the absence of other PCAT-encoding genes in the genome, we hypothesized that PctA and PctB form an alternate T1SS that can secrete sufficient levels of sulfated RaxX to activate XA21.

### PctB partially compensates for the loss of RaxB

To assess the potential role of PctB in RaxX maturation and secretion, we deleted *raxB* and *pctB* singly or together from the PXO99 genome. The resultant mutant strains (designated Δ*raxB*, Δ*pctB*, and Δ*raxB* Δ*pctB*, respectively) were inoculated onto rice plants by clipping leaf tips with scissors dipped in bacterial suspension. Lesion lengths and bacterial densities *in planta* measured 14 days after inoculation were comparable to wildtype PXO99 on Taipei 309 (TP309) rice plants, which lack *Xa21* (Figures 5A and 5C; Song et al., 1995). This indicates that deletion of genes encoding these putative transporters do not directly compromise *Xoo* virulence under the tested conditions.

**Figure 5.**
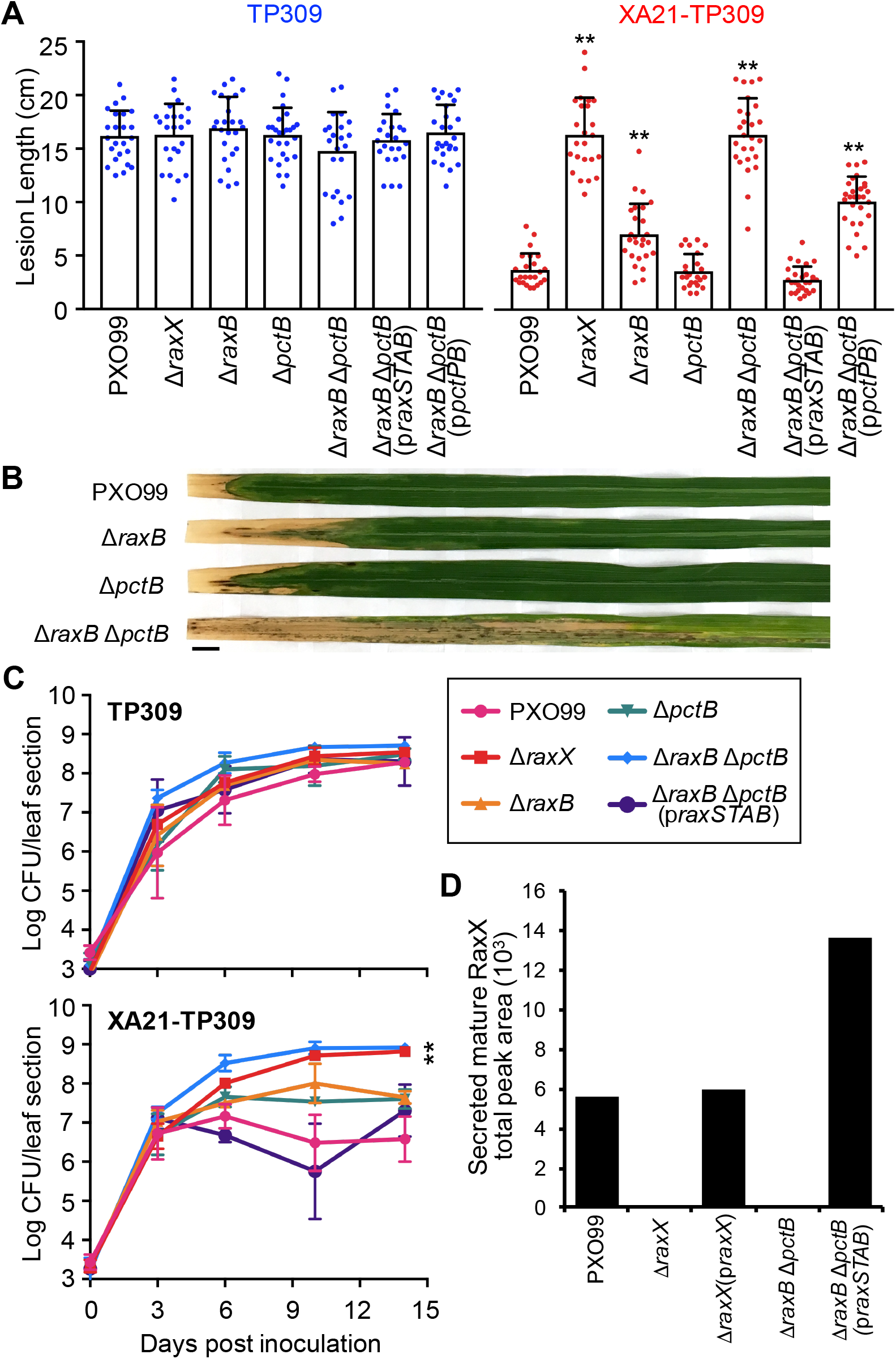
The Δ*raxB* Δ*pctB* double mutant does not secrete RaxX, but secretion can be restored with the p*raxSTAB* plasmid. (A-C) Rice plants were inoculated by scissor clipping with PXO99-derived strains. (A) Bars represent the mean + SD of lesions (cm) measured 14 days post inoculation (dpi) of TP309 (blue dots) or XA21-TP309 (red dots) plants (n = 21-28). (B) Inoculated XA21-TP309 leaves 14 dpi. Scale bar, 1 cm. (C) Bacterial densities in planta. Data points represent the mean log CFU per 10-cm leaf section ± SE (n = 3). (D) SRM-MS was used to detect the presence of mature RaxX in *Xoo* supernatants. Bars represent the total peak area of the chromatograms shown in Figure S3C. **p < 0.01; compared to PXO99 using Dunnett’s test. Similar results were observed in 5 (A) or 2 (C) other experiments, except Δ*raxB* formed lesions comparable to PXO99 in roughly half. See also Figures S2 and S3.

We next tested the effect of the mutant strains on XA21-TP309 rice, a derivative of TP309 that expresses *Xa21* (Song et al., 1995), to assess XA21-dependent responses. As previously reported, XA21-TP309 rice were resistant to PXO99 infection and had short lesions (<5 cm) and low bacterial densities (6.8 × 10^6^ CFU/leaf) *in planta* (Figures 5A-5C; Song et al., 1995). We observed that the Δ*raxB* mutant formed short (< 5 cm) to intermediate (6-9 cm) lesions on XA21-TP309, which varied between experiments, but the Δ*pctB* mutant consistently formed short (<5 cm) lesions similar to PXO99 (Figures 5A and 5B). Both Δ*raxB* and Δ*pctB* single mutants accumulated to populations of ~4.5 x 10^7^ CFU/leaf, roughly 6.6-fold higher than PXO99 (Figure 5C). In contrast, the Δ*raxB* Δ*pctB* double mutant formed long lesions (>14 cm) and accumulated to 8.4 x 10^8^ CFU/leaf, over 4- and 120-fold higher than the PXO99 population, respectively (Figures 5A-5C). These phenotypes were comparable to the Δ*raxX* control (Figures 5A and 5C). Similarly, the Δ*raxA* Δ*pctA* double mutant, containing mutations in genes for the predicted periplasmic adaptor proteins RaxA and PctA, formed longer lesions than the Δ*raxA* and Δ*pctA* single mutants (Figure S2). Collectively, these results suggest that sulfated RaxX is not secreted in both the Δ*raxA* Δ*pctA* and Δ*raxB* Δ*pctB* double mutants, allowing these strains to evade XA21. These results also suggest that PctA and PctB can partially compensate for the loss of RaxA and RaxB, respectively.

### RaxB is the primary PCAT required for RaxX maturation and secretion

The Δ*raxB* Δ*pctB* double mutant accumulates to high population levels in XA21-TP309 plants (Figure 5C), indicating that this strain can evade detection by the XA21 immune receptor. This result supports the hypothesis that these PCATs secrete sulfated RaxX. Given that *raxB* is genetically clustered with *raxX* (Figure 4A), we hypothesized that RaxB functions as the primary transporter of RaxX and is sufficient to activate XA21 in the absence of PctB. To test this hypothesis, we complemented the Δ*raxB* Δ*pctB* double mutant with p*raxSTAB*, a broad host range vector expressing the entire *raxSTAB* operon with the presumptive native promoter. On XA21-TP309, the p*raxSTAB* plasmid restored *raxB* gene expression as well as activation of XA21-mediated immunity, resulting in short lesions and bacterial densities comparable to wildtype PXO99 (Figures S3A, 5A and 5C). Similarly, p*raxSTAB* restored the ability of the Δ*raxA* Δ*pctA* double mutant to activate XA21-mediated immunity (Figure S2). These results suggest that the RaxAB system is sufficient to secrete RaxX and activate XA21 in the absence of the PctAB system.

As a control, we also expressed p*raxSTAB* in the wildtype PXO99 background. We observed no significant alterations on lesion development with the addition of p*raxSTAB* (Figure S2), suggesting that overexpression of *raxSTAB* did not directly contribute to enhanced activation of the XA21-mediated immune response.

We next assessed, using SRM-MS, if mature RaxX was secreted in the Δ*raxB* Δ*pctB* double mutant. RaxX was detected in the supernatant of the Δ*raxB* Δ*pctB* double mutant only in the presence of the p*raxSTAB* plasmid (Figure 5D). We detected SRM transition peaks of mature RaxX in the p*raxSTAB*-containing strain at the same retention time as wildtype PXO99 and the Δ*raxX* mutant complemented with plasmid-expressed *raxX* (p*raxX*) controls, which were collected and analyzed in the same batch (Figure S3B). Together, these results indicate that RaxB is a necessary and sufficient PCAT for RaxX maturation and secretion.

We also tested if PctB is sufficient to restore RaxX secretion in the Δ*raxB* Δ*pctB* double mutant. For this purpose, we transformed the Δ*raxB* Δ*pctB* double mutant with a broad host range vector expressing the *pctPB* cluster with its presumptive native promoter (p*pctPB*). The p*pctPB* plasmid restored *pctB* gene expression but failed to fully restore XA21-mediated recognition of the Δ*raxB* Δ*pctB* double mutant (Figures S3A and 5A). The p*pctPB*-complemented Δ*raxB* Δ*pctB* double mutant still formed intermediate-to-long lesions (~10 cm) roughly 3-fold longer than PXO99 (Figure 5A). This result suggests that PctB alone cannot effectively secrete RaxX to activate XA21-mediated immunity.

### RaxB and PctB carry conserved residues characteristic of bifunctional PCATs

PCATs such as *E. coli* CvaB function as both a transporter and a protease (Håvarstein et al., 1995; Wu and Tai, 2004). The peptidase domain of this class of ABC transporters contains 2 conserved motifs: a cysteine (C) motif and histidine (H) motif (Figure 6A; Dirix et al., 2004; Håvarstein et al., 1995). We found that both RaxB and PctB contain sequences that match the conserved C- and H-motifs, including the conserved Cys, His, and Asp residues (Cys-28/12, His-101/85, Asp-117/101 of RaxB/PctB, respectively) predicted to form the active site catalytic triad (Figure 6A; Wu and Tai, 2004; Wu et al., 2012). Additionally, they contain the conserved Gln residue (Gln-22/6 of RaxB/PctB, respectively) proposed to form the oxyanion hole that stabilizes the Cys-His ion pair (Figure 6A; Ménard et al., 1991; Wu et al., 2012). In contrast, ABC transporters that have a degenerate C39-like domain (CLD) with no proteolytic activity, such as the *E. coli* hemolysin transporter HlyB, do not contain the catalytically essential Cys or conserved Gln (Figure 6A; Lecher et al., 2012). The presence of the complete Cys-His-Asp/Asn catalytic triad and conserved oxyanion hole Gln characteristic of bifunctional PCATs is consistent with the hypothesis that RaxB and PctB also possess proteolytic activity to process RaxX.

**Figure 6.**
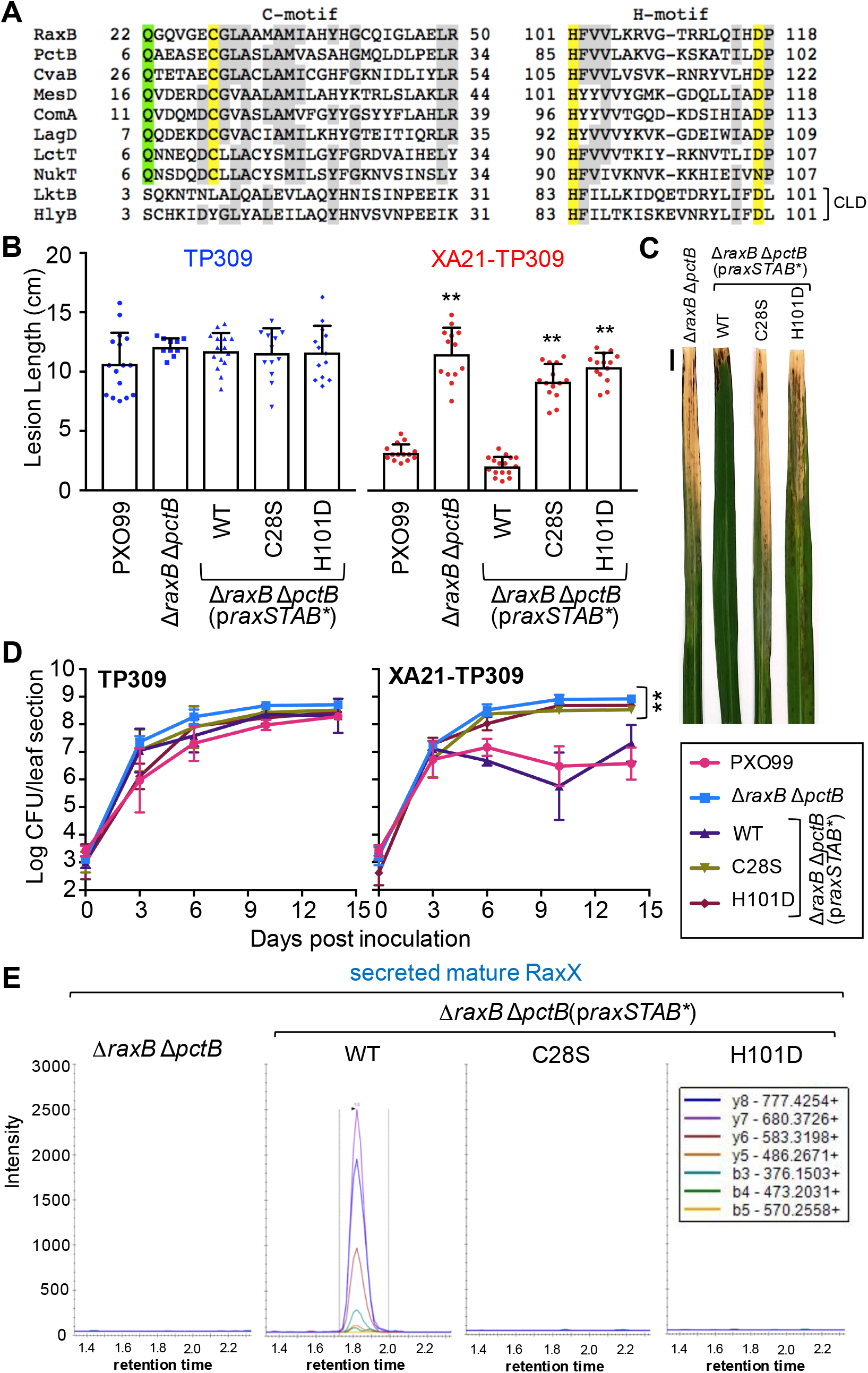
Mutation of the RaxB peptidase catalytic triad impairs RaxX maturation and secretion. (A) Alignment of the peptidase C- and H-motifs of RaxB and PctB to select PCATs or ABC transporters with a C39-like domain (CLD; listed in method details). Cys-His-Asp/Asn catalytic triad highlighted in yellow, oxyanion hole Gln in green, and residues common in at least 6 of the transporters in gray. (B-D) Rice plants were inoculated by scissor clipping with the indicated strains. (B) Bars represent the mean + SD of lesion measurements (cm) on TP309 (blue dots) or XA21-TP309 (red dots) plants 14 dpi (n = 10-16). (C) Inoculated XA21-TP309 leaves 14 dpi. Scale bar, 1 cm. (D) Bacterial densities in planta. Data points represent the mean log CFU per 10-cm leaf section ± SE (n = 3). Performed at the same time as Figure 5C. (E) SRM-MS chromatograms of mature RaxX tryptic peptide detected in supernatants from the Δ*raxB* Δ*pctB* double mutant-derived strains. **p < 0.01; compared to PXO99 using Dunnett’s test. Similar results were observed in 5 (B) or 2 (D) other independent experiments.

### Mutation of the RaxB peptidase domain catalytic triad impairs RaxX maturation and secretion

We next assessed the role of the RaxB peptidase domain in RaxX maturation. For these experiments, we transformed the Δ*raxB* Δ*pctB* double mutant with a derivative of p*raxSTAB* containing site-directed missense substitutions of the active-site cysteine (Cys-28) or histidine (His-101) of the RaxB peptidase domain. Cys-28 was mutated to Ser (C28S) and His-101 was mutated to Asp (H101D), which we hypothesized would abolish proteolytic activity by disrupting formation of the predicted Cys-28, His-101, Asp-117 catalytic triad as demonstrated with *E. coli* CvaB (Wu and Tai, 2004).

Expression of the C28S and H101D mutant derivatives impaired p*raxSTAB*-complemention of the Δ*raxB* Δ*pctB* double mutant phenotype on XA21-TP309 but had no affect on the TP309 control plants (Figures 6B-6D). Wildtype p*raxSTAB* fully complemented the Δ*raxB* Δ*pctB* double mutant phenotype on XA21-TP309, resulting in short lesions (<4 cm) and low bacterial densities (3.9 × 10^7^ CFU/leaf). In contrast, expression of the mutated peptidase derivatives of p*raxSTAB* resulted in long lesions (>9 cm) and high bacterial densities (10^8^ CFU/leaf) comparable to the strain without plasmid (Figures 6B-6D). These results indicate that the predicted RaxB peptidase domain is catalytically active and necessary to process and secrete biologically active RaxX.

To assess if mature RaxX is secreted into supernatants of the C28S or H101D RaxB peptidase mutants, we carried out SRM-MS analysis. We detected high SRM transition peaks corresponding to mature RaxX in supernatants of the Δ*raxB* Δ*pctB* double mutant strain carrying wildtype p*raxSTAB* (Figure 6E). In contrast, SRM transitions corresponding to mature RaxX were not detected in the Δ*raxB* Δ*pctB* double mutant carrying p*raxSTAB* with either the C28S or H101D mutation (Figure 6E). We also failed to detect SRM transitions corresponding to mature RaxX in the Δ*raxX* negative control and the Δ*raxB* Δ*pctB* double mutant without plasmid (Figures S3B and 6E). The lack of detectable secreted RaxX in the RaxB peptidase mutants supports our inoculation results, providing further evidence that RaxB serves as the RaxX maturation protease.

### Mutation of the Gly-Gly cleavage site compromises RaxX maturation and secretion

The RiPP precursor typically carries a cleavable N-terminal leader that is removed by a protease (Figure 1A). PCATs cleave the N-terminal leader of substrates downstream of a GG-motif, which contains Gly-Gly or Gly-Ala immediately preceding the cleavage site (Håvarstein et al., 1994, 1995). Based on the observation that key residues in the RaxB catalytic peptidase domain are required for RaxX secretion (Figure 6), we hypothesized that RaxX contains an N-terminal GG-motif that is removed by RaxB. In support of this hypothesis, our SRM-MS analysis indicated that the N-terminal 39 residues of RaxX are removed prior to secretion (Figure 3). We noted that this leader peptide ends with three conserved glycines and contains conserved hydrophobic residues at positions −4, −7, −12, and −15 distal to the cleavage site, a pattern typical of PCAT substrates and necessary for interaction with the peptidase (Figures 3A, 7A and S4A; Aucher et al., 2005; Kotake et al., 2008). These observations suggest that the predicted RaxX leader resembles the leader of GG-motif-containing peptides typically processed and secreted by PCATs.

**Figure 7.**
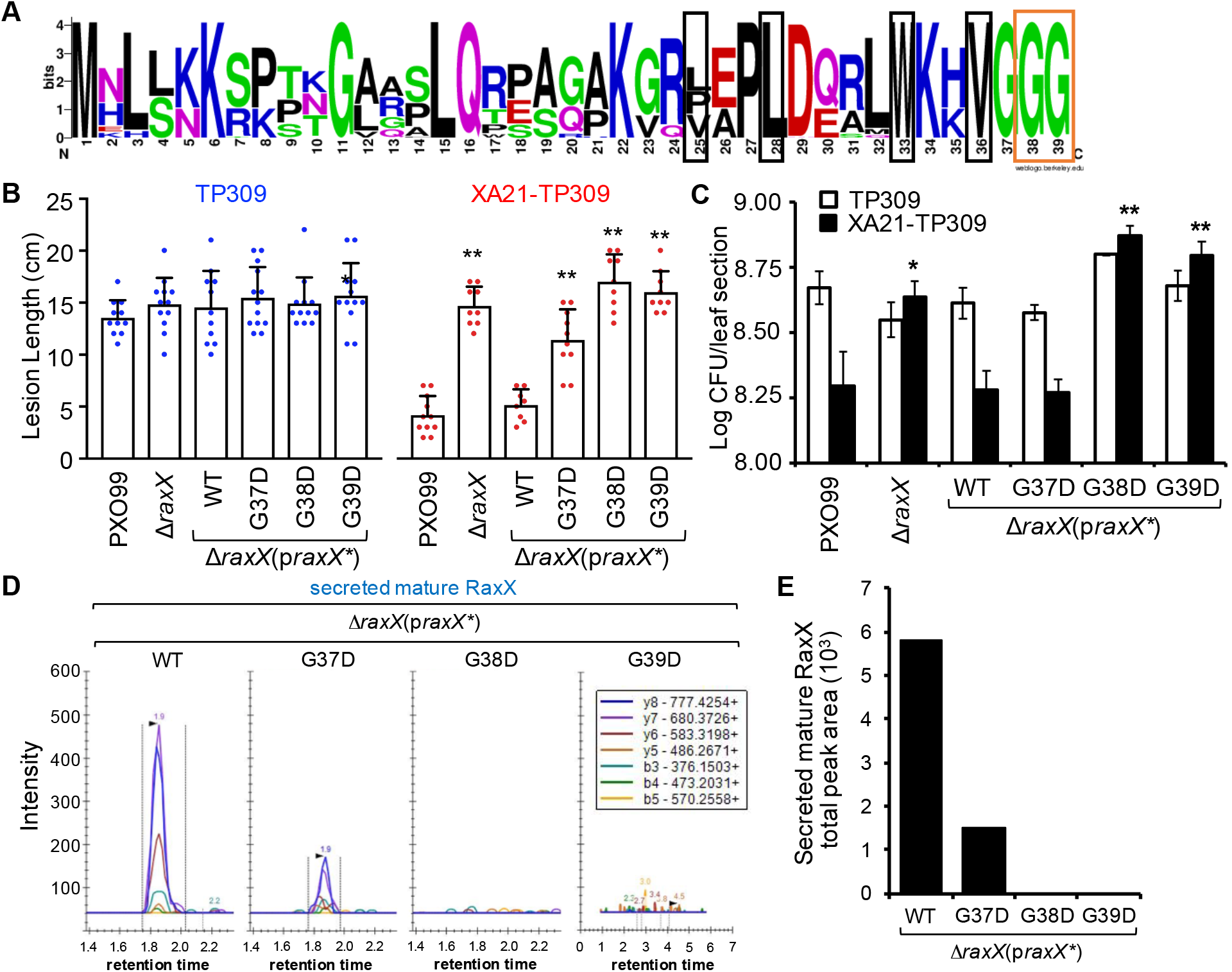
Mutation of the Gly-Gly cleavage site compromises RaxX maturation and secretion. (A) Sequence logo of the leader from all *Xanthomonas* RaxX alleles. Conserved Gly-Gly preceding the cleavage site boxed in orange; hydrophobic residues in conserved positions typical of PCAT substrates boxed in black. (B-C) Rice plants were inoculated by scissor clipping. (B) Bars represent the mean + SD of lesion measurements (cm) on TP309 (blue dots) or XA21-TP309 (red dots) plants 14 dpi (n = 8−13). Similar results were observed in at least 6 independent experiments. (C) Bars represent the mean log CFU/leaf section ± SE measured 14 dpi (n = 4). (D) SRM-MS chromatograms of mature RaxX tryptic peptide detected in *Xoo* supernatants. (E) Quantification of the total peak area shown in D. *p < 0.05; **p < 0.01; compared to PXO99 using Dunnett’s test. See also Figure S4.

Studies of the peptide mesentericin Y105 (MesY) produced by *Leuconostoc mesenteroides* indicate that the GG-motif preceding the MesY cleavage site is critical for efficient processing and secretion (Aucher et al., 2005). Based on this observation, we hypothesized that the GG-motif in the predicted RaxX leader sequence is critical for RaxB-mediated processing of RaxX. To test this hypothesis, we generated site-directed missense substitutions of the conserved glycine residues (37-39) preceding the cleavage site of RaxX. As Aucher et al. (2005) demonstrated that secretion of MesY was significantly reduced or completely abolished when Gly at the −2 position from the cleavage site was mutated to Ala or Asp or when Gly at the −1 position from the cleavage site was mutated to Arg or Asp, we generated homologous missense substitutions of RaxX Gly-(37-39). As a control, we also generated missense substitutions of Gly-Ala pairs further upstream of the cleavage site: Gly-11, Ala-12 and Gly-20, Ala-21. These mutant variants of RaxX were expressed as derivatives of the p*raxX* plasmid in the Δ*raxX* mutant background.

The resulting mutants were inoculated onto rice plants. These mutants formed long (>12 cm) lesions comparable to PXO99 on the TP309 control plants (Figures 7B, S4B, and S4C). On XA21-TP309, Δ*raxX* formed long lesions (>15 cm) and this phenotype was complemented to short lesions by p*raxX* as previously reported (Figures 7B, S4B, and S4C; Pruitt et al., 2015). Similarly, p*raxX* derivatives containing mutations in RaxX residues Gly-11, Ala-12, Gly-20, or Ala-21 still complemented the Δ*raxX* mutant phenotype (Figures S4B and S4C). This result suggests that these residues are not necessary for processing nor secretion of RaxX. In contrast, p*raxX* derivatives containing mutations in Gly-(37-39) were impaired in their ability to complement the Δ*raxX* mutant phenotype (Figures 7B, S4B, and S4C). Expression of the G37D and G38A derivatives resulted in intermediate lesions (11-12 cm) but the G38D, G38R, and G39D derivatives resulted in long lesions (>15 cm) similar to the Δ*raxX* mutant without plasmid control (Figures 7B, S4B, and S4C). Bacterial densities were also comparable among the Δ*raxX* mutant and the G38D and G39D-expressing strains (Figure 7C). These results suggest that Gly-(37-39), which immediately precede the cleavage site, are essential for RaxX maturation and/or secretion.

We next assessed if we could detect secreted RaxX in supernatants of *Xoo* strains carrying mutations in the leader peptide. We did not detect SRM transitions of RaxX above background levels in supernatants of the Δ*raxX* mutant carrying the G38D and G39D derivatives (Figure 7D), suggesting that RaxX is neither processed nor secreted when Gly-38 or Gly-39 is mutated to Asp. In contrast, we detected RaxX in supernatants of the Δ*raxX* mutant carrying the G37D p*raxX* derivative although at reduced levels compared to expression of wildtype p*raxX* (1.5 × 10^3^ vs. 5.8 × 10^3^ total peak area, respectively; Figure 7E). The accumulation of intermediate amounts of RaxX peptide secreted from the G37D mutant strain is consistent with the formation of intermediate lesion lengths on XA21-TP309 plants inoculated with this strain (Figures 7B and 7E). Taken together, our results suggest that proper RaxX maturation and secretion, which require the Gly-38, Gly-39 cleavage site, are necessary steps in the biogenesis of the immunogenic peptide.

## DISCUSSION

### Tyrosine sulfation of RaxX enhances XA21 binding

RaxX immunogenicity is dependent on sulfation of Tyr-41 (Pruitt et al., 2015). Our MST results demonstrate that this modification increases the affinity of RaxX binding to XA21^ECD^ (Figure 2), resulting in a K_D_ comparable to the RaxX concentration needed to trigger the half maximal immune response of XA21 (~20 nM) and induce root growth (Pruitt et al., 2015, 2017). Tyrosine sulfation has similarly been observed to enhance interaction of the disulfated pentapeptide phytosulfokine (PSK) with its cognate receptor PSKR1 in *Arabidopsis* and carrot. Based on crystal structure studies, the two sulfate groups on PSK form hydrogen bonds and van der Waals packing with residues on PSKR1 (Wang et al., 2015). This sulfate group-mediated interaction stabilizes PSKR1, which in turn allosterically induces heterodimerization with the SERK (somatic embryogenesis receptor-like kinase) co-receptor, resulting in activation of PSKR1 (Wang et al., 2015). Analogous structural studies will provide insights into how sulfation of RaxX Tyr-41 contributes to the RaxX-XA21 interaction.

### RaxX shares similarities to RiPPs

RaxX is predicted to mimic the activity of the plant peptide hormone PSY1 (Pruitt et al., 2017). Our results indicate that RaxX is ribosomally synthesized as a precursor that is proteolytically processed by RaxB into a mature peptide (Figure 1B). This mature RaxX peptide includes the same residues covered by RaxX16, which is sufficient to activate XA21-mediated immunity and promote root growth in rice (Figure 3A; Pruitt et al., 2017). These residues also precisely overlap the conserved 13-residue region found in mature PSY1 from *Arabidopsis* and in putative PSY-like proteins from various plant species including rice (Amano et al., 2007; Pruitt et al., 2017). Furthermore, the mature RaxX and PSY1 peptides start with the same residues, Asp-Tyr, with the critical sulfated tyrosine at the same position. Mature RaxX therefore intimately resembles host PSY-like proteins.

The small size (<10 kDa), presence of a post-translational modification (sulfated tyrosine in this case), and proteolytic maturation of RaxX and PSY1 are defining characteristics of RiPPs (Arnison et al., 2013). In addition to PSY1, plants produce other secreted tyrosine-sulfated peptides that regulate plant growth including PSK, root meristem growth factor (RFG), and Casparian strip integrity factor (CIF) (Matsubayashi and Sakagami, 1996; Matsuzaki et al., 2010; Doblas et al., 2017; Nakayama et al., 2017). Similar to RaxX and PSY1, PSK, RFG, and CIF are proteolytically processed from larger precursor peptides and secreted. We therefore propose that these peptides be classified as RiPPs.

RiPPs are subdivided into over 20 groups according to their biosynthetic machinery and structural features imparted primarily by the post-translational modification(s) that decorate the core (Arnison et al., 2013). However, more groups continue to be added as new RiPPs are identified. Given the similarities of RaxX, PSY1, PSK, RFG, and CIF, we propose that these peptides be classified as a new group of RiPPs defined by the presence of the sulfated tyrosine.

### Does PctB process and secrete other RiPPs?

Our results indicate that RaxAB is the cognate RaxX maturation and secretion system and can be partially complemented by PctAB. This functional redundancy is rather unusual given the low sequence identity shared between the two systems, which are also independently distributed among Xanthomonads. For instance, the *X. albilineans* genome contains *pctAPB* but not *raxX* nor *raxSTAB*, whereas the *X. euvesicatoria* genome contains *raxX* and *raxSTAB* but not *pctAPB*. These observations suggest that RaxX is not the cognate substrate for PctB.

Because PctB is predicted to have a functional C39 peptidase domain (Figure 6A), we hypothesize that its cognate substrate is a GG-motif-containing peptide. Based on the annotated PXO99 sequence (NC_010717.2), two hypothetical genes, PXO_RS14845 and PXO_RS14850 (here referred to as *pct-92* and *pct-75*, respectively), are encoded upstream of *pctA* (Figure 4B). *pct-92* and *pct-75* are predicted to encode 92- and 75-residue peptides, respectively, that contain a GG-like leader of 28 and 19 amino acids, respectively (Figure S4A). If Pct-92 and Pct-75 are both PctB substrates, then this would suggest that PctB can tolerate leaders that vary in sequence and length. Such substrate tolerance has been observed in PCATs such as LagD, HalT, and LicT from *Lactococcus lactis*, *Bacillus halodurans*, and *B. licheniformis*, respectively, which process and transport two cognate peptides that function together as a two-component antimicrobial molecule (Caetano et al., 2014; Håvarstein et al., 1995; Lawton et al., 2007). Because the leader of the two peptide components can vary in sequence and length (Figure S4A), these transporters exhibit a relaxed specificity and sometimes tolerate noncognate substrates (Caetano et al., 2014). This may explain why PctB is able to tolerate RaxX, at least to some extent. However, we have yet to confirm that *pct-92* and *pct-75* encode actual peptides. It is also unclear if these putative peptides are post-translationally modified as no known modification enzymes are encoded in the same genomic region (Figure 4B). Therefore, further studies are needed to analyze these putative PctB substrates and identify essential determinants that allow PctB to distinguish substrates.

### Does the RaxX leader peptide have functional roles in RiPP biosynthesis other than peptidase recognition?

In the RaxX leader peptide, residues proximal to the peptidase cleavage site are well conserved in *raxX* alleles from all *Xanthomonas* species (Figure 7A). This includes Gly-38 and Gly-39 preceding the cleavage site, which are critical for proper RaxX maturation and secretion (Figure 6), as well as the hydrophobic residues that occupy positions −4, −7, and −12 from the cleavage site (Val-36, Trp-33, and Leu-28, respectively), which are proposed to position the Gly-Gly cleavage site in the enzyme’s active site (Ishii et al., 2010). These features are also conserved in leaders of PCAT substrates, including RiPPs and unmodified peptides (Figure S4A), suggesting that this region of the RaxX leader is important for recognition by the RaxB and PctB peptidase domain. However, it is unclear if sulfation of RaxX, which occurs one residue from the cleavage site, is required for recognition by RaxB or possibly impairs recognition by PctB.

Because many unmodified peptides have a shorter leader peptide, the extended N-terminus of RaxX may have other functional roles aside from maturation and secretion that we have yet to identify.

One proposed role of the leader peptide is to keep the precursor inactive during biosynthesis as demonstrated for *Lactococcus lactis* lacticin 481, an antimicrobial RiPP of the lanthipeptide class (Xie et al., 2004). However, the observation that exogenous application of RaxX60, representing the full-length precursor peptide, is able to activate XA21-dependent immune responses (Pruitt et al., 2015) and promote root growth in *Arabidopsis* and rice (unpublished data), does not support a role for the RaxX leader in maintenance of an inactive state.

Another possibility is that the leader peptide serves as a recognition motif for post-translational modification enzymes. We previously detected sulfated non-processed RaxX in *Xoo* cell lysates (Pruitt et al., 2015), suggesting that sulfation occurs while the leader is attached. However, under *in vitro* conditions, RaxST can sulfate a peptide derivative of human CCR5, containing only an aspartate preceding the tyrosine (Han et al., 2012). This result suggests that the RaxX leader is not required for RaxST function. Similarly, the lacticin 481 precursor LctA, which is post-translationally modified by dehydration and cyclization reactions catalyzed by LctM, does not require the leader for its modification (Levengood et al., 2007). Instead, the presence of the leader peptide promotes the efficiency of LctM modification of the LctA core peptide even when supplemented *in trans* as a synthetic peptide (Levengood et al., 2007) or covalently fused to LctM (Oman et al., 2012). Further studies are needed to determine if the RaxX leader helps recruit RaxST or enhance its sulfotransferase activity in an analogous manner.

### Knowledge of RiPP biosynthesis can be used to engineer a diversity of tyrosine sulfated molecules

*raxX* is genetically clustered with *raxST*, *raxA* and *raxB*, which are each involved in its biosynthesis. The simplicity of the RaxX biosynthetic machinery makes it an ideal system to study how sulfation affects substrate specificity and activity of the biosynthetic proteins. This information can provide insights into the development of strategies to engineer new tyrosine sulfated molecules such as hybrid RiPP products. Burkhart et al. (2017) used such an approach to demonstrate that sequences recognized by unrelated RiPP modification enzymes, including thiazoline-forming cyclodehydratases and lanthipeptide synthetases, can be combined into a single chimeric leader within a precursor peptide. Co-expression of this chimeric construct with the appropriate modification enzymes led to the production of hybrid RiPPs *in vivo*. With the addition of leader-independent “tailoring” modification enzymes, the chemical diversity of these hybrid RiPPs can be further extended. We envision that the Rax system can be similarly exploited to increase the chemical diversity and potency of sulfated molecules that may be useful as therapeutic agents to suppress infection or activate the immune response of plants and animals.

### Conclusion

Here we show that PCAT processing is required for both the maturation and export of bacterial RaxX, an immunogen that binds a eukaryotic receptor. These results suggest that RaxX is a tyrosine sulfated RiPP, a group not previously described. We further identified and defined the RaxX leader, which contains residues that are critical for its maturation and secretion. These results set the stage for leader peptide-guided biosynthetic strategies to extend the chemical diversity of hybrid RiPPs.

## MATERIALS AND METHODS

### Peptides

Tyrosine sulfated (sY) and/or nonsulfated (nY) versions of RaxX13 (DYPPPGANPKHDP), RaxX16 (DYPPPGANPKHDPPPR), and RaxX21 (HVGGGDYPPPGANPKHDPPPR) were synthesized. The synthetic peptides were ordered from Pacific Immunology (Ramona, CA), with the exception of RaxX16, which was ordered from Peptide 2.0 (Chantilly, VA). Tyrosine sulfated PSY1 was synthesized by the Protein Chemistry Facility at the Gregor Mendel Institute. All of the peptides were resuspended in ddH_2_0. Recombinant full-length RaxX (RaxX60-sY and RaxX60-nY) was isolated from an engineered *E. coli* strain using an expanded genetic code approach as previously described (Pruitt et al., 2015).

### XA21^ECD^ expression and purification

The extracellular domain (ECD) of XA21 (residues 23 - 649) was amplified using primers listed in Table S2 and inserted into the baculovirus transfer vector pMeIBac B1 (Invitrogen, Carlsbad, CA) using RecA-mediated Sequence and Ligation Independent Cloning (SLIC) strategy (Scholz et al., 2013). A C-terminal Strep II-9xHis tag was fused to XA21^ECD^ and verified by Sanger Sequencing. XA21^ECD^-StrepII-9xHis was produced by secreted expression in baculovirus-infected High Five insect cells, harvested 72 hours post-infection. Subsequently, the protein was affinity purified by Ni-NTA chromatography (Ni Sepharose excel, GE Healthcare, Chicago, IL) and subjected to size-exclusion chromatography column (Superdex 200 16/60, GE Healthcare) pre-equilibrated with 50 mM NaH_2_PO_4_/Na_2_HPO4 pH 7.5, 200 mM NaCl, 5% Glycerol. All of the purification steps were checked by SDS-PAGE.

### Microscale thermophoresis (MST)

Purified XA21^ECD^ was labelled with a fluorescent dye using Monolith™ Protein Labelling Kit RED-NHS (Amine Reactive; NanoTemper Technologies, San Francisco, CA). Fluorescently-labelled XA21^ECD^ (at constant concentration 0.0133 μM) was mixed with varying peptide concentrations (ranging from 0.00006 to 2 μM) in buffer containing 50 mM NaH_2_PO4/Na_2_HPO4 pH 7.5, 200 mM NaCl, 5% Glycerol and 0.001% Tween. Approximately 4-6 μL of each sample was loaded in a fused silica capillary (NanoTemper Technologies). Measurements were performed at room temperature in a Monolith NT.115 instrument at a constant LED power of 65% and MST power of 60%. Measurements were performed repeatedly on independent protein preparations to ensure reproducibility. The data were analysed by plotting peptide concentrations against percent changes of normalized fluorescence. Curve fitting was performed using PALMIST software (Scheuermann et al., 2016).

### Bacterial strains and culture

A list of bacterial strains used in this study are provided in Table S1. *Escherichia coli* DH5α, which was used for general cloning, was cultured in lysogeny broth (LB) with the appropriate antibiotics at 37°C (shaking at 230 rpm for liquid cultures). *Xoo* strains were routinely cultured at 28°C on peptone sucrose agar (PSA) with the appropriate antibiotics unless otherwise indicated. Nutrient broth or agar was initially used to generate the mutant strains. When needed, sucrose was added to a final concentration of 5% and antibiotics were added. For proteomic analysis, cells were grown on XOM2 (Tsuge et al., 2002) agar, a plant-mimicking medium. Cephalexin was used at 20 μg/mL, kanamycin at 50 μg/mL, and geneticin at 50 μg/mL.

### Protein extraction

*Xoo* proteins were prepared from strains grown on XOM2 agar plates, which was found to induce *raxX* expression. After incubation at 28°C for 2 days, the fully-grown cells were scraped from the plates and re-suspended in 30 mL of water to loosen the extracellular polysaccharide matrix containing the secretome. The cells were pelleted by centrifugation (12k rpm, 45 min, 4°C). The supernatants, which contained the secreted proteins, were harvested and concentrated to 0.5-1 mL using Amicon^®^ Ultra Centrifugal Filters Ultracel^®^-3K (Millipore, Burlington, MA, cat#UFC900324). Total protein was also extracted from the cell pellet. The cells were resuspended and lysed by sonication in lysis buffer (25mM Tris pH 8, 300mM NaCl, 1mM PMSF, 2.5 mM EDTA, and 1mg/ml lysozyme). Soluble proteins were separated by centrifugation (12k rpm, 45 min, 4°C).

All of the samples for mass spectrometry were digested with 1 μg of trypsin (Promega, Madison, WI) in the presence of 5 mM DTT. The tryptic peptides were desalted with a C18 column (Harvard apparatus, Holliston, MA, cat#74-4601) and eluted with 80 % acetonitrile, 0.1 % formic acid.

### LC-SRM-MS Analysis

The SRM targeted proteomic assays were performed, as described previously (Batth et al., 2014), on an Agilent 6460 QQQ mass spectrometer system coupled with an Agilent 1290 UHPLC system (Agilent Technologies, Santa Clara, CA). Peptides were separated on an Ascentis Express Peptide C18 column [2.7-mm particle size, 160-Å pore size, 5-cm length × 2.1-mm inside diameter (ID), coupled to a 5-mm × 2.1-mm ID guard column with same particle and pore size, operating at 60°C; Sigma-Aldrich, St. Louis, MO] operated at a flow rate of 0.4 ml/min via the following gradient: initial conditions were 95% solvent A (0.1% formic acid), 5% solvent B (99.9% acetonitrile, 0.1% formic acid). Solvent B was increased to 35% over 5 min, and was then increased to 80% over 0.2 min, and held for 2 min at a flow rate of 0.6 mL/min, followed by a ramp back down to 5% B over 0.5 min where it was held for 1.5 min to re-equilibrate the column to original conditions. The eluted peptides were ionized via an Agilent Jet Stream ESI source operating in positive ion mode with the following source parameters: gas temperature = 250°C, gas flow = 13 liters/min, nebulizer pressure = 35 psi, sheath gas temperature = 250°C, sheath gas flow = 11 liters/min, capillary voltage = 3500 V, nozzle voltage = 0 V. The data were acquired using Agilent MassHunter version B.08.02. Acquired SRM data were processed by using Skyline software version 3.70 (MacCoss Lab Software, Seattle, WA).

### Immunoblotting

The proteins were separated by SDS-PAGE, transferred to PVDF membranes (Bio Rad, Hercules, CA cat#162-0177) by electrophoresis and then analyzed by immunoblot using standard procedures. RaxX was detected using an anti-RaxX antibody that was raised in rabbit against RaxX11 (PPGANPKHDPP), derived from residues 43-53 of the 60-residue RaxX precursor (GenScript, Piscataway, NJ). Alkaline phosphatase conjugated anti-rabbit IgG (Sigma, cat#A3687) was used as secondary antibody and CDP-*Star* (Roche, Basel, Switzerland, cat#12041677001) was used as substrate. The anti-RaxX and anti-rabbit IgG antibodies were used at dilutions of 1:2000 and 1:20,000, respectively. The membranes were exposed to X-ray films or visualized using the Gel-Doc XR^+^ system (Bio-Rad).

### Identification of the *pct* locus

PctB (PXO_RS14825) was identified by NCBI Protein BLAST analysis of the PXO99 genome (NC_010717.2) using the RaxB (PXO_RS06015) N-terminal 150 residues as query under the default settings. PctP (PXO_RS14830) and PctA (PXO_RS14840) were identified by algorithms from NCBI in the rhomboid family (CDD:304416) and the type_I_hylD (CDD:330454) domain classification, respectively. Pct-92 (PXO_RS14845) and Pct-75 (PXO_RS14850) were annotated as hypothetical proteins.

### *Xoo* mutation and complementation

*Xoo* mutants were generated in the Philippine race 6 strain PXO99^A^ (referred to as PXO99 in this manuscript;Hopkins et al., 1992). The mutant strain Δ*raxX* and the complemented strain Δ*raxX*(p*raxX*) used in this study were previously reported (Pruitt et al., 2015). The rest of the mutant strains: Δ*raxB*, Δ*pctB*, Δ*raxB* Δ*pctB*, Δ*raxA*, Δ*pctA*, and Δ*raxA* Δ*pctA*, were generated by double homologous recombination using the pUFR80 suicide vector (Castañeda et al., 2005). Briefly, two flanking sequences upstream and downstream of the target gene were independently PCR-amplified with their respective primer sets A, B and C, D (Table S2) using Phusion High-Fidelity DNA polymerase (Thermo Scientific, Waltham, MA). Both PCR products were fused together by overlap extension PCR (Horton et al., 1989) using external primers A and D. This PCR product was then cloned into pUFR80 using the sites EcoRI/HindIII or BamHI/HindIII. The resultant plasmids, which were verified by sequencing, were electroporated into PXO99 (Δ*raxB* or Δ*raxA* was used instead for generation of the Δ*raxB* Δ*pctB* or Δ*raxA* Δ*pctA* double mutant, respectively). The electroporated cells were then plated onto nutrient agar (NA) with kanamycin (50 μg/mL) and grown at 28°C for 5-6 days. Positive integrants were screened by PCR, grown in nutrient broth without kanamycin overnight, and then plated onto NA containing 5% sucrose to select for the second crossover event. Colonies were analyzed by PCR to identify deletion mutants and verified by sequencing the PCR product.

For complementation, all of the plasmids (listed in Table S3) were constructed using standard recombinant DNA techniques using the primers listed in Table S2. p*raxX-ha* was generated by cloning the *raxX* coding sequence into pENTR/D-TOPO (Invitrogen, Carlsbad, CA), which was then recombined into pLN615, a gateway destination vector containing an HA tag (Guo et al., 2009). p*raxSTAB* and p*pctPB* were derived from pVSP61 (Loper and Lindow, 1994), into which the indicated gene clusters and 0.4 kb of the upstream sequence, presumed to contain the native promoter, were cloned. Expression of *raxSTAB* and *pctPB* were chosen instead of independent expression of *raxB* and *pctB*, respectively, to account for potential translational coupling with *raxA* and *pctP*, respectively. Site-directed mutagenesis of catalytic residues of the RaxB peptidase domain and residues of the RaxX leader were generated in p*raxSTAB* and p*raxX*, respectively. All of the plasmids were verified by sequencing. The plasmids were transformed into *Xoo* competent cells by electroporation, plated onto PSA with the appropriate antibiotic, and grown at 28°C. Colonies were analyzed by PCR and verified by sequencing the PCR product.

### Rice plants

The *Oryza sativa* ssp. *japonica* rice variety TP309 and a transgenic TP309 line derived from I106-17 (XA21-TP309), which carries the *Xa21* gene driven by its own promoter (Song et al., 1995) were used for rice inoculations. Native TP309 does not contain *Xa21*. TP309 and XA21-TP309 rice plants were grown as previously described (Pruitt et al., 2015). Briefly, seeds were germinated in distilled water at 28°C for 1 week then transplanted to sandy soil in 5.5-inch square pots (3 seedlings per pot). Plants were grown in tubs filled with fertilizer water in a greenhouse. Six-week old plants were then transferred to a growth chamber at least 2 days prior to inoculation. The growth chamber was set to 28°C/24°C, 80%/85% humidity, and 14/10-hour lighting for the day/night cycle.

### Rice inoculation

*Xoo* strains were cultured on PSA plates with appropriate antibiotics at 28°C. Bacteria from the plates were resuspended in sterile water at a density of 10^8^ colony-forming units (CFU)/mL and then inoculated onto rice plants using the scissor clipping method (Kauffman, 1973). Briefly, surgical scissors were dipped in bacterial suspension and used to clip the tips of the two uppermost expanded leaves in each tiller, representing one biological replicate. The length (cm) of the water-soaked lesions extending from the site of inoculation were measured 14 days after inoculation. The lesions of at least 8 biological replicates, each a mean of the lesions formed on the two leaves, were measured. At least 3 independent experiments were performed with similar results unless indicated otherwise.

Analysis of bacterial growth *in planta* was performed as previously described (Bahar et al., 2014) with minor modifications. Briefly, leaves (10-cm section from the site of inoculation) were harvested for each sample set at each time point. The two leaf sections from each tiller, representing one biological replicate, were cut to 2 mm pieces and incubated in sterile water for 2 hours at 28°C shaking at 230 rpm. This suspension was serially diluted, plated onto PSA plates with cephalexin (20 μg/mL), and incubated at 28°C. Colonies were counted 2 days later and the cell number per 10-cm leaf section was calculated. Three to four biological replicates, each a mean number of colonies from 4 technical replicates, were measured for each data point. Three independent experiments were performed with similar results.

For *in planta* bacterial gene expression analysis, TP309 leaves were harvested 6 days after inoculation. The leaves were cut 4 cm from the site of inoculation and the 4-cm section from two leaves of each tiller, representing one biological replicate, were snap-frozen in liquid nitrogen and processed as described below.

### RNA Extraction and RT-qPCR

Total RNA was extracted from homogenized rice leaf tissue using TRIzol reagent (Invitrogen) following the manufacturer’s instructions and then treated with the TURBO DNA-free kit (Ambion) to remove residual genomic DNA. cDNA was synthesized using the High-Capacity cDNA Reverse Transcription Kit (Applied Biosystems, Foster City, CA) according to the manufacturer’s instructions. Quantitative PCR (qPCR) was performed on a Bio-Rad CFX96 Real-Time System coupled to a C1000 Thermal Cycler (Bio-Rad) using the iTaq Universal SYBR Green Supermix (Bio-Rad) and 100 ng of cDNA. The primers used are listed in Table S2. The 2^(−ΔΔCt) method was used to calculate the relative changes in gene expression normalized to the *ampC2* reference gene control and the PXO99 wildtype strain. Three biological replicates, each a mean of two technical replicates, were analyzed. Three independent experiments were performed with similar results.

### Sequence analysis and visualization

The C- and H-motifs of the RaxB and PctB peptidase domains were identified from alignments of the peptidase domain of the *E. coli* colicin V transporter CvaB (CAA40744.1; Wu and Tai, 2004), the *Leuconostoc mesenteroides* mesentericin B105 and Y105 transporter MesD (CAA57402.1; Aucher et al., 2005; Morisset and Frère, 2002), the *Streptococcus pneumoniae* competence-stimulating peptide transporter ComA (AAA69510.1; Ishii et al., 2006), the *Lactococcus lactis* lactococcin G transporter LagD (ACR43772.1; Håvarstein et al., 1995), the *Lactococcus lactis* lacticin 481 transporter LctT (AAC72259.1; Furgerson Ihnken et al., 2008), and the *Staphylococcus warneri* nukacin ISK-1 transporter NukT (NP_940774.2; Nishie et al., 2009), as well as the peptidase-like domain of the *Aggregatibacter actinomycetemcomitans* leukotoxin transporter LktB (CAA37906.1; Guthmiller et al., 1995), and the *E. coli* alpha-hemolysin transporter HlyB (KKA61973.1; Lecher et al., 2012). The alignment was performed using the default settings in Clustal Omega (European Bioinformatics Institute; EMBL-EBI; Sievers et al., 2011).

The sequence logo of the RaxX leader was constructed using WebLogo (Crooks et al., 2004) with the predicted leader sequence of RaxX alleles from various *Xanthomonas* species identified by Pruitt et al. (2017). The RaxX leader was aligned with the predicted leader of the candidate PctB substrates Pct-92 and Pct-75 as well as the experimentally verified leader sequences from the *E. coli* colicin V precursor CvaC (AAX22078.1; Håvarstein et al., 1994), the *Leuconostoc mesenteroides* mesentericin B105 and Y105 precursors MesB and MesY (AAD54223.1 and AAP37395.1, respectively; Morisset and Frère, 2002; Aucher et al., 2005); the *Streptococcus pneumoniae* competence-stimulating peptide precursor ComC (AAC44895.1; Ishii et al., 2006), the *Lactococcus lactis* lactococcin alpha and beta precursors LagA and LagB, respectively (ACR43769.1 and ACR43770.1, respectively; Håvarstein et al., 1995), the *Lactococcus lactis* lacticin 481 precursor LctA (AAC72257.1; Furgerson Ihnken et al., 2008), and the *Staphylococcus warneri* nukacin ISK-1 precursor NukA (NP_940772.1; Nishie et al., 2009).

### Statistical analysis

Statistical analysis was performed using JMP Pro 13 (SAS Institute Inc., Cary, NC) or GraphPad Prism 7 (GraphPad Software, La Jolla, CA) and specified in figure legends. Differences were considered to be significant when p < 0.05.

## ACKNOWLEDGMENTS

This work was supported by NIH GM59962, NIH GM122968, and NSF PGRP grant IOS-1237975. This work was also partially supported by the Office of Biological and Environmental Research’s Genomic 275 Science program within the U.S. Department of Energy Office of Science, under award number DE-AC02-05CH11231.

## AUTHOR CONTRIBUTIONS

D.D.L., A.J., O.B., C.P., Y.B., and P.C.R. conceived and designed the research. D.D.L., A.J., Y.C., K.P., R.P., L.C., K.L., and C.A. performed experiments. D.D.L., A.J., Y.C., K.P., and V.S. analyzed data. D.D.L., A.J., V.S., and P.C.R. wrote the manuscript with assistance and feedback from the other authors.

